# UKBCC: a cohort curation package for UK Biobank

**DOI:** 10.1101/2020.07.12.199810

**Authors:** Isabell Kiral, Nathalie Willems, Benjamin Goudey

## Abstract

**Summary:** The UK Biobank (UKB) has quickly become a critical resource for researchers conducting a wide-range of biomedical studies (Bycroft *et al.,* 2018). The database is constructed from heterogeneous data sources, employs several different encoding schemes, and is disparately distributed throughout UKB servers. Consequently, querying these data remains complicated, making it difficult to quickly identify participants who meet a given set of criteria. We have developed UK Biobank Cohort Curator (UKBCC), a Python tool that allows researchers to rapidly construct cohorts based on a set of search terms. Here, we describe the UKBCC implementation, critical sub-modules and functions, and outline its usage through an example use case for replicable cohort creation.

**Availability:** UKBCC is available through PyPi (https://pypi.org/project/ukbcc) and as open source code on GitHub (https://github.com/tool-bin/ukbcc).

## 1 Introduction

The UK Biobank (UKB) is a rich, open-access, and continuously growing database that has become a key resource for medical and genomics research (Bycroft *et al*. (2018)). The four main sources of data (self-reported fields, clinical and biological measurements, in-patient hospital records, and primary care data are split across two files. Since the coding of phenotypes differs across these sources, incorrectly mapping between codes, joining databases, or only querying a subset of relevant datafields can lead to reduced sample size or missing exclusionary criteria when constructing a cohort for scientific study.

Some tools have been developed that attempt to simplify interaction with the UKB. BiobankRead (Schneider-Luftman and Crum, 2019) provides a command-line interface and searches record-level data, ukbtools (Hanscombe *et al*., 2019) allows keyword searches and computes some summary statistics, and ukbRest (Pividori and Im, 2018) offers a REST Api to interface with a local copy of UKB data. However, there exist no tool that provides a comprehensive abstraction layer from the complex encoding schemes within UKB or supports multiple criteria for cohort construction.

UKBCC addresses these common pain points when dealing with UKB, including searching both main and record-level databases, automatically saving cohorts as well as search criteria, and supporting complex queries through a simple interface.

## 2 Implementation

Figure 1 provides an overview of the key modules implemented within UKBCC, outlining their function when creating a cohort from a set of search terms. The package has two modes of interaction: 1) ‘interactive’ mode, which uses a command-line interface to automate most of the steps with minimal human input (highlighted in yellow), and 2) ‘import’ mode, in which the various modules and functions can be imported into existing scripts (blue boxes).

**Fig. 1.**
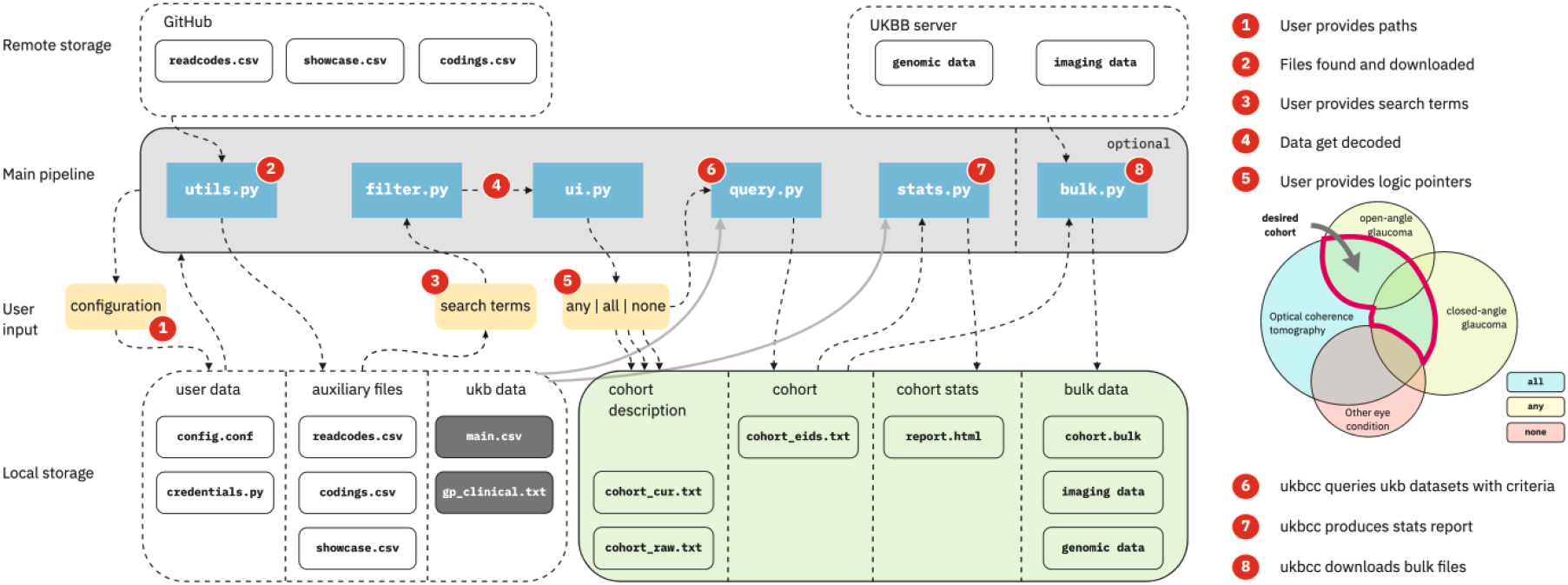
UKBCC pipeline. The main file orchestrates user input and data usage. The package modules are depicted in blue, user input in yellow, and created output files are shown in the bottom right. Step 1) Provide paths of necessary files and output directories in a configuration file, which, unless otherwise specified, will be created interactively (‘interactive’ mode). Step 2) The package downloads auxiliary files used for the decoding of different data sources and stores the files locally (paths specified in step 1). Note that this step can be skipped for future runs. Step 3) Search terms are provided in plain text. Step 4) The package scans datafield descriptions and codes for all search terms. Step 5) Logic pointers are provided for the returned datafield:code combinations. In this example. Step 6) UKB data is scanned using the cohort description. Logic operations are carried out automatically and the package returns the set of participant IDs that are stored locally. Step 7) The package creates a summary statistics report. Step 8) Bulk data is downloaded if applicable.

We will illustrate the typical process of cohort construction through an example use case of studying glaucoma. The input search terms will include ‘glaucoma’, ‘eye condition’, and ‘optical coherence tomography’ (OCT). The aim is to establish a cohort of participants that have **‘any’** of two conditions: ‘open-angle glaucoma’ and/or ‘closed-angle glaucoma’, where **‘all’** selected patients must have OCT data and **‘none’** of the participants will have a non-glaucoma eye condition.

In order to facilitate the use of plain-text search terms, the UKBCC module automatically consolidates all encoding schemes within UKB (Read 2/3, ICD9/10 and main datafield encodings), and returns a single, searchable data structure containing the human-readable forms of the codes (filter module). This step leverages several files to construct the final data structure, translating the encoded datafields and values in the main dataset, as well as the the record-level data which uses a mixture of standard and custom encoding schemes. This step is crucial, as it simultaneously abstracts two levels of the complex mapping procedure. The returned data structure is then searched for matches against the keywords. In our use case, the keywords return 461 relevant fields. This number alone illustrates the breadth of relevant diagnoses available within UKB. At this stage, the user can easily select the relevant datafields using the command-line interface. In our example, the user would only select datafields pertaining to ‘open-angled glaucoma’, ‘closed-angle glaucoma’, other eye conditions, and ‘optical coherence tomography’, resulting in a subset of 20 datafields.

A second challenge lies in supporting both inclusion and exclusion criteria within a single query or set of search terms. UKBCC addresses this by performing conditional logic after the selection of decoded datafield:values. This step is broken down into 3 selection criteria: **‘any’**, **‘all’**, and **‘none’**. Returning to our example use case, the user would select ‘none’ for all datafield:value combinations that relate to non-glaucoma eye conditions, they would select ‘all’ for the datafield:value combinations for OCT data, and ‘any’ for the datafield:value combinations relating to glaucome sub-types. The Venn diagram shown in Figure 1 illustrates the set of desired participants. Once these conditions are set, the package will search the available databases for individuals adhering to the selection criteria, and return a list of participant IDs. This file, as well as the criteria for the cohort, are saved to a user-specified location, enabling both cohort replication and rapid adjustments of existing cohorts. All functionality is modular, and thus can be imported into existing scripts separately as well (‘import’ mode).

## 3 Discussion

Whilst a resource such as the UKB presents a great opportunity for detailed scientific study, it has a significant entry barrier due to its complexities. Tools such as UKBCC aim to lower this barrier by implementing a user-friendly package that relies on keyword search terms to construct cohorts. Table 1 provides an overview of the key functionalities of UKBCC compared to existing tools. UKBCC addresses both key limitations of existing tools, as well as consolidates a comprehensive set of functionalities within a single framework. Importantly, the package enables less computationally advanced research groups to access the wealth of information within UKB by automating most of the steps required to construct cohorts, while existing tools require more advanced technical knowledge (e.g experience with docker containers, postgreSQL databases, etc.).

**Table 1.** Comparison of UKBCC functionality against related tools.

| Functionality | BiobankRead | ukbtools | ukbREST | UKBCC |
| --- | --- | --- | --- | --- |
| Command-line interface | Yes | No | No | Yes |
| Keyword search | No | Yes | No | Yes |
| Searches record-level data | Yes | No | No | Yes |
| Conditional logic | Yes | Yes | No | Yes |
| Cohort replication | No | No | Yes | Yes |
| Abstraction of encoding | No | No | No | Yes |
| Provides API | No | Yes | No | Yes |
| Generates statistics | No | Yes | No | Yes |
| Automates file downloads | No | No | No | Yes |

The first version of the UKBCC tool has some limitations that we expect to address in future iterations. UKB contains a number of different record-level data tables. Currently, UKBCC’s support is limited to the GP clinical SQL table. Furthermore, the flexibility of user interaction through the command line interface is limited. We intend to provide a web portal to further facilitate interaction with the package.

## References

Bycroft, C. et al. (2018). The UK Biobank resource with deep phenotyping and genomic data. Nature, 562(7726), 203–209.

Hanscombe, K. B. et al. (2019). ukbtools: An R package to manage and query UK Biobank data. PLOS ONE, 14(5), e0214311.

Pividori, M. and Im, H. K. (2018). ukbREST: efficient and streamlined data access for reproducible research in large biobanks. Bioinformatics, 35(11), 1971–1973.

Schneider-Luftman, D. and Crum, W. R. (2019). BioBankRead: Data pre-processing in Python for UKBiobank clinical data. bioRxiv, page 569715.

